# Cytoplasm localized ARID1B promotes oncogenesis in pancreatic cancer by activating RAF-ERK signaling

**DOI:** 10.1101/830075

**Authors:** Srinivas Animireddy, Padmavathi Kavadipula, Viswakalyan Kotapalli, Swarnalata Gowrishankar, Satish Rao, Murali Dharan Bashyam

## Abstract

The ARID1B/BAF250b subunit of the human SWI/SNF chromatin remodeling complex is a canonical nuclear tumor suppressor. Immunohistochemistry on a pancreatic cancer tissue microarray revealed significant ARID1B cytoplasmic localization that correlated with advanced tumor stage and lymph node positivity. Identification of the nuclear localization signal (NLS) using *in silico* prediction and subcellular localization studies facilitated evaluation of a possible cytoplasmic function for ARID1B. A cytoplasm-restricted ARID1B-NLS mutant was significantly compromised to regulate transcription activation and tumor suppression functions, as expected. Surprisingly however, cytoplasm-localized ARID1B could bind c-RAF and PPP1CA causing stimulation of RAS-RAF-ERK signaling and β-catenin transcription activity in pancreatic cancer cells. More importantly, cytoplasmic ARID1B resulted in an induction of cell growth and migration in pancreatic cancer cell lines that was dependent on ERK signaling and caused increased tumorigenesis in nude mice. NLS peptides representing mutations identified from pancreatic cancer samples exhibiting ARID1B cytoplasmic localization or curated from cancer somatic mutation database were significantly compromised to effect nuclear localization of a reporter protein. ARID1B cytoplasmic localization correlated significantly with active forms of ERK and β-catenin in primary pancreatic tumor samples. ARID1B may therefore promote oncogenesis through non-canonical cytoplasm-based gain of function mechanisms in addition to dysregulation in the nucleus.

## Introduction

<u>SWI</u>tch/<u>S</u>ucrose <u>N</u>on-<u>F</u>ermentable (SWI/SNF) complex, originally discovered in yeast, functions as the primary chromatin remodeler in humans during ontogeny and adult life ^7^. It is a large (∼2 MDa) evolutionarily conserved multi-functional complex that uses energy from ATP hydrolysis to make nucleosomal DNA accessible for various regulatory proteins facilitating nuclear processes such as transcription, DNA repair and maintenance of chromosomal stability ^25^. The multi subunit complex includes one of two ATPase subunits BRG1 or BRM, three core components namely INI1, BAF155 (<u>B</u>RG1 <u>A</u>ssociated <u>F</u>actor) and BAF170, four mutually exclusive DNA binding subunits namely the <u>A</u>T-<u>r</u>ich interaction <u>d</u>omain 1A (ARID1A or BAF250a), ARID1B (BAF250b)^33^, ARID2 (BAF180)^34^ or GLTSCR1/GLTSCR1L^21^, in addition to up to ten varying accessory subunits. Inactivating mutations in genes encoding one of several SWI/SNF subunits are identified in up to 20% of cancers ^12^ attesting to the importance of this complex in tumor suppression.

ARID1B is a ubiquitously expressed subunit of the SWI/SNF complex ^8^. ARID1B levels increase during differentiation of embryonic stem cells ^13^ while its ectopic expression induces *TP53*/*CDKN1A* causing cell cycle arrest in HeLa cells ^9^ suggesting a possible tumor suppressor function. ARID1B loss of function events including mutations ^6^, chromosomal rearrangements ^26^ and DNA methylation ^14^ are reported in several cancer types. In our previous study, we described a tumor suppressor role for nuclear ARID1B in pancreatic cancer (PaCa) ^14^. Here, we report cytoplasmic localization of ARID1B in several pancreatic tumor samples that correlate significantly with advanced tumor stage and lymph node positivity. Surprisingly, an NLS-mutant version of ARID1B localized to the cytoplasm and stimulated RAF-ERK signaling and β-catenin transcription activity causing increased tumorigenic features in pancreatic cancer cell lines as well as in primary tumors.

## Results

### ARID1B exhibits cytoplasmic localization in PaCa

We previously reported loss of ARID1B expression in a significant proportion of PaCa samples by immunohistochemistry (IHC) on a tissue microarray (TMA) ^14^. We have now confirmed the results on a larger TMA (figures 1a and S1; Table S1). A careful analysis surprisingly revealed cytoplasmic localization of ARID1B in a significant proportion of tumor (but not normal) samples confirmed independently by two pathologists blinded for the study (figures 1a-b; Table S1). IHC performed on another nuclear protein (p53) did not reveal cytoplasmic stain in any sample (data not shown) indicating that ARID1B cytoplasmic localization was not an artefact. More importantly, ARID1B cytoplasmic localization correlated significantly with aggressive clinical features including advanced tumor stage and lymph node positivity (figure 1c). We therefore proceeded to test a possible role for cytoplasm localized ARID1B in PaCa progression.

**Figure 1.**
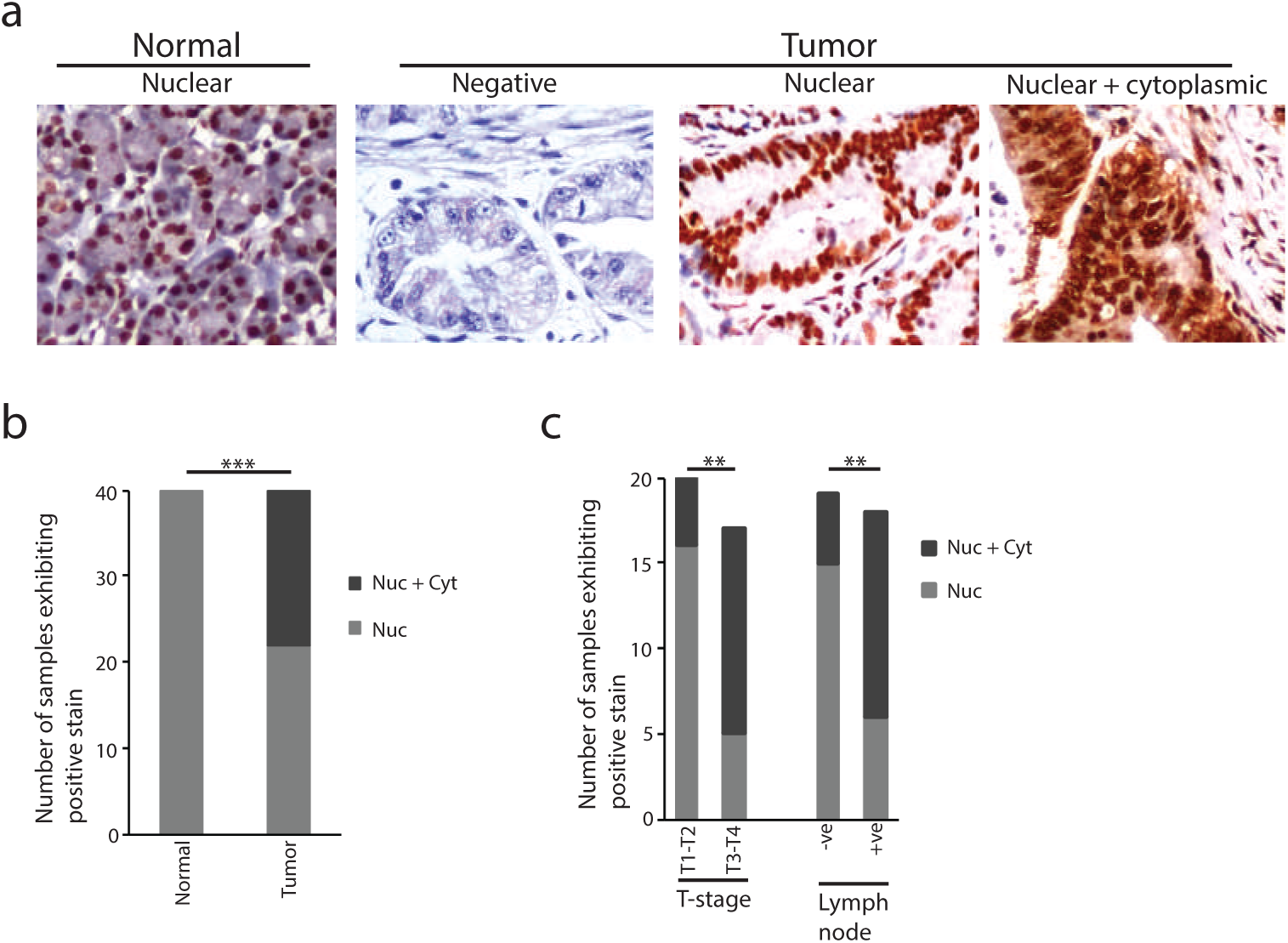
Identification of ARID1B aberrant cytoplasmic localization in PaCa samples. **a,** IHC based evaluation of ARID1B intracellular localization in pancreatic primary tumors. **b,** Analysis of number of samples exhibiting ARID1B nuclear vs nuclear plus cytoplasmic stain in tumor and normal samples (forty tumor/normal pairs). **c,** Association of nuclear vs nuclear plus cytoplasmic stain with tumor stage or lymph node status (thirty seven tumor samples). ARID1B negative samples as well as chronic pancreatitis, groove pancreatitis, cystic adenoma, and gastro-intestinal stromal tumor samples were excluded from the analyses depicted in panels b and c. Differences were considered significant at a p value less than 0.05 (*, p<0.05; **, p<0.01; ***, p<0.001). Nuc, nuclear staining; Nuc + Cyt, nuclear plus cytoplasmic staining; T1-T2, low tumor stage; T3-T4, advanced tumor stage; -ve, negative; +ve, positive.

### ARID1B possesses a classical bipartite NLS sequence

In order to evaluate the significance of its cytoplasmic localization in pancreatic tumor samples we decided to determine the effect of restricting ARID1B to the cytoplasm in PaCa cell lines. Since this can be achieved by simply inactivating its NLS, we proceeded to define the ARID1B NLS.

Based on a comparison of subcellular localization exhibited by several truncation constructs, the probable location of the ARID1B NLS appeared to be between the ARID and the BAF250_C domains (figure 2a; grey box). A computational search using cNLS mapper (http://nls-mapper.iab.keio.ac.jp) revealed several putative NLS sequences of which the one with the highest score (Table S2) localized within the region (figure 2b) predicted by the truncation constructs. This putative twenty one amino acid sequence appeared to belong to the classical bipartite NLS family that is defined by two clusters of basic amino acids (Lysine-Arginine in this case) separated by a spacer region of 9-12 amino acids (10 in this case). Sequence alignment of human ARID1B (Uniprot ID: Q8NFD5) with its orthologues from 41 metazoan species (figure S2a) followed by WebLogo based identification of the consensus (figure 2b) revealed significant conservation of the predicted NLS. The ARID1B NLS was also significantly similar to that of its paralog ARID1A (figure S2b).

**Figure 2.**
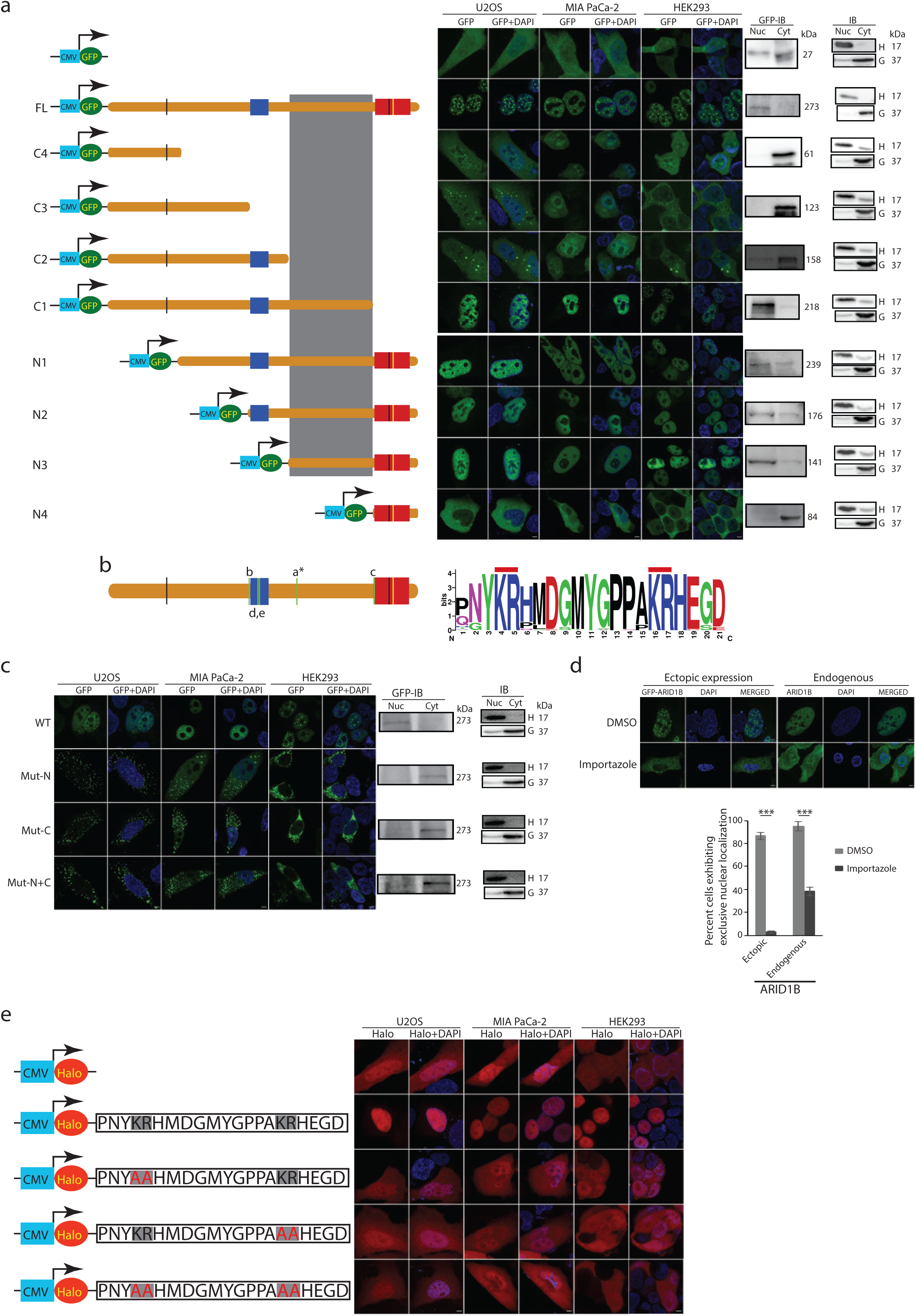
Identification and characterization of the ARID1B NLS. **a,** ARID1B truncations identify the putative region harboring the NLS. Diagrammatic depiction of constructs expressing GFP alone or fused with full length or various ARID1B truncations generated in pDEST-N-EGFP is shown on the left. The ARID1B full length construct is designated as FL and each C- and N-terminal deletion construct is labelled as C1-C4 and N1-N4, respectively. ARID1B domains are shown in various colors: blue, ARID domain; red, BAF250_C domain; black, LXXLL motif; yellow, B/C box motif. A rectangular gray box depicts probable location of the NLS. Intracellular localization of each GFP-tagged protein was detected in three different cell lines (indicated) using GFP fluorescence (shown in the middle) and by immunoblotting performed separately for the nuclear and cytoplasmic fractions (only in HEK293; shown on the right). Immunoblotting for Histone H3 (labelled as ‘H’) and GAPDH (labelled as ‘G’) were used as controls for nuclear and cytoplasmic fractions, respectively (shown on the far right). Nuc, nuclear fraction; Cyt, cytoplasmic fraction; IB, immunoblotting. **b,** Left panel depicts location of five putative ARID1B NLS sequences (shown as thin green vertical bars and labelled a-e) identified using cNLS mapper. ‘*’ indicates position of the NLS sequence selected for characterization. The right panel depicts the consensus ARID1B NLS sequence derived using WebLogo (weblogo.berkeley.edu) with default settings based on the alignment of ARID1B amino acid sequences from 42 vertebrate species (enumerated in legend to figure S2a) using Clustal Omega. Each of the two highly conserved basic amino acid clusters (Lysine-Arginine) are indicated by a red line on the top. **c,** Representative results from transfected cells via GFP fluorescence as well as from GFP immunoblotting performed separately on nuclear and cytoplasmic fractions (only in HEK293). WT, full length ARID1B; Mut-N, full length ARID1B harboring mutated N terminal basic amino acid cluster (AA instead of KR); Mut-C, full length ARID1B harboring mutated C terminal basic amino acid cluster (AA instead of KR); Mut-N+C, full length ARID1B harboring mutated N and C terminal basic amino acid clusters; IB, immunoblotting; Nuc, nuclear fraction; Cyt, cytoplasmic fraction; H, Histone H3 immunoblotting as a positive control for the nuclear fraction; G, GAPDH immunoblotting as a positive control for the cytoplasmic fraction. **d,** Evaluation of ARID1B intracellular localization following Importazole treatment on U20S cells using GFP fluorescence and ARID1B antibody based immunofluorescence analyses. The graph depicts results from fluorescence quantitation. **e,** ARID1B NLS peptide alone can translocate a fused reporter (Halo tag) into the nucleus. Diagrammatic representation of each NLS peptide variant expression construct is shown on the left (the basic amino acid clusters are indicated by grey shading and the mutant residues are indicated in red color) and fluorescence signal from representative cells expressing the particular variant is shown on the right. Scale bars are 10µm for all fluorescence images. Differences were considered significant at a p value less than 0.05 (***, p<0.001).

Alanine substitutions in either one or both basic amino acid clusters caused ARID1B to be significantly redistributed to the cytoplasm (figure 2c). Nuclear localization achieved through the classical (mono or bipartite) NLS is regulated by the importin α/β pathway^16^. Importazole (an importin β blocker^29^) treatment caused a significant reduction in nuclear localization of ectopically expressed as well as endogenous ARID1B (figures 2d and S3a) providing further evidence towards the authenticity of the predicted NLS. Further, NLS peptides harboring Alanine substitutions in either or both of the two basic amino acid clusters lost the ability of nuclear translocation exhibited by the wild type NLS peptide (figures 2e and S3b). These results suggest that the predicted NLS was not only necessary but perhaps sufficient to translocate an unrelated protein into the nucleus.

### Similar to a null mutant, cytoplasm-localized ARID1B loses tumor suppressor and transcription regulatory functions

Ectopic expression of the cytoplasm-localized version of ARID1B (mut-N+C from figure 2c) severely compromised senescence induction when compared to wild type ARID1B in the PaCa cell line MIA PaCa-2 (figure 3a). Similarly, the ARID1B-NLS mutant was significantly restricted in its ability to transcriptionally activate *CDKN1A* (p21) and *TP53* in both MIA PaCa-2 and PANC-1 cells (figures 3b-c), corroborated further using *CDKN1A* promoter-luciferase assay (figure S4a). We also tested induction of *CDKN1B*/p27, a well-established inducer of senescence not previously shown to be regulated by ARID1B or the SWI/SNF complex. Interestingly, wild type ARID1B caused significant induction of *CDKN1B*/p27 at transcript and protein levels (figures 3b-c) confirmed further using promoter-luciferase assay (figure S4a). The cytoplasmic form of ARID1B was however unable to induce *CDKN1B*/p27 (figures 3b-c and S4a).

**Figure 3.**
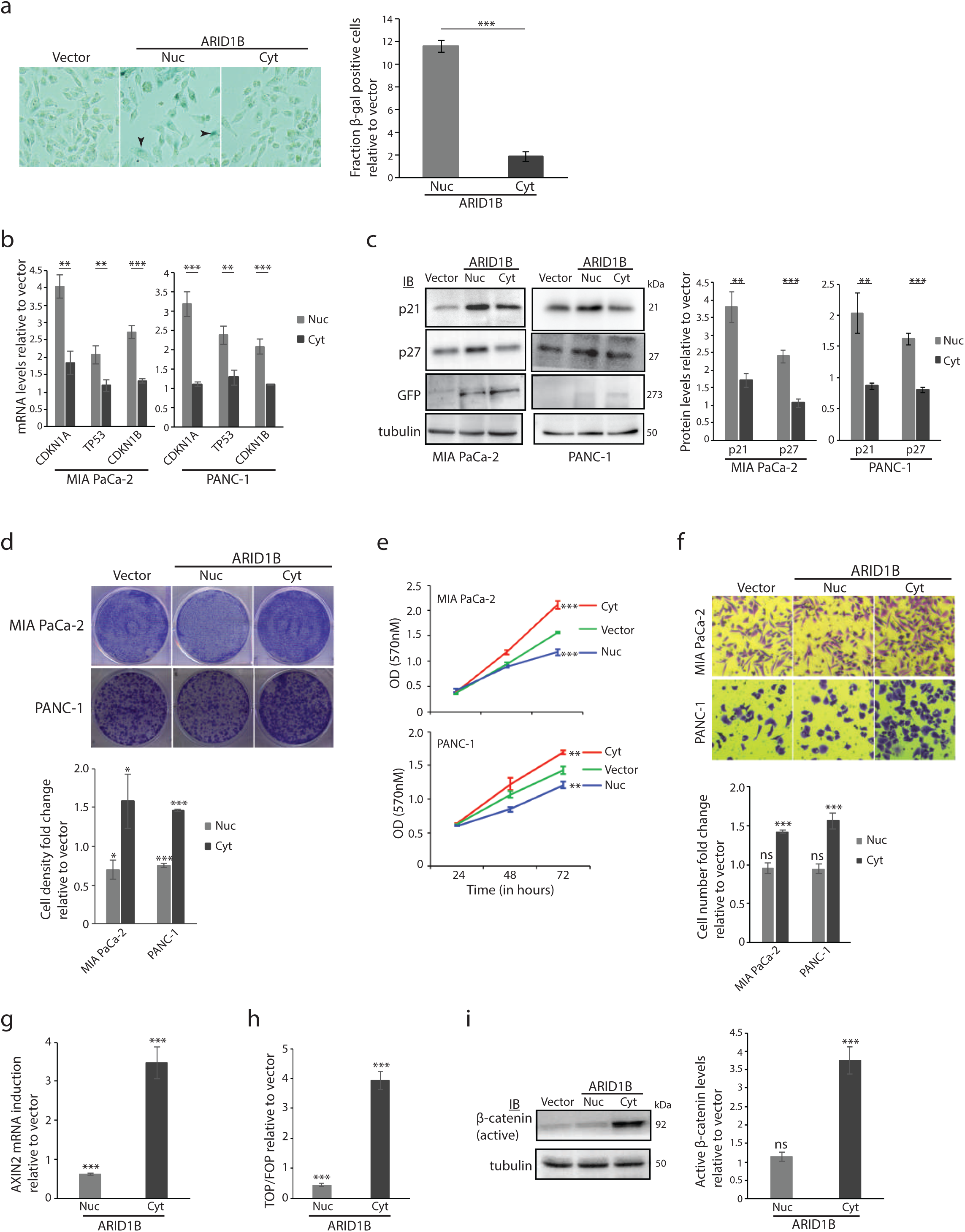
Cytoplasm restricted ARID1B is compromised for transcription regulatory and tumor suppressor functions but promotes oncogenic properties. **a,** Senescence-associated β-galactosidase assay. Left panel shows representative images of β-galactosidase stain while the right panel shows quantitation. β-galactosidase positive cells are indicated by black arrowheads. **b-c,** ARID1B NLS mutant is compromised to induce transcription of canonical tumor suppressor targets *CDKN1A*, *TP53* as well as *CDKN1B* measured at both transcript (b) and protein (c) levels. **d-f,** Ectopic expression of cytoplasmic version of ARID1B results in increased tumorigenic potential of pancreatic cancer cells as measured by crystal violet staining (d), MTT (e) and transwell migration (f) assays. **g-i,** Cytoplasm-localized ARID1B elevates β-catenin transcription activity in MIA PaCa-2 cells as evaluated by measuring *AXIN2* transcript levels (g) and relative promoter activity of TOPFlash over FOPFlash (h) as well as by quantitation of active (non-phosphorylated at Ser45) β-catenin levels (i). IB, immunoblotting. Differences were considered significant at a p value less than 0.05 (*, p<0.05; **, p<0.01; ***, p<0.001; ns, not significant). Nuc, nuclear (wild type) ARID1B; Cyt, cytoplasmic (NLS mutant) ARID1B.

### Unlike a null mutant, cytoplasm-localized ARID1B exhibits oncogenic functions

We confirmed ability of wild type ARID1B to cause reduction in cell viability and growth ^9, 30^ in PaCa cells using crystal violet staining and MTT assay (figures 3d-e). Surprisingly however, the cytoplasmic form of ARID1B appeared to exhibit a neo-morphic function causing a significant increase in cell viability in the same assays (figures 3d-e). Further, cytoplasm localization of ARID1B also caused an increased migratory potential in cells, while there was no effect upon expression of the wild type (nuclear) form (figure 3f).

Given that wild type (nuclear) ARID1B inhibits β-catenin nuclear function ^31^, we evaluated status of transcriptional activation mediated by β-catenin upon ectopic expression of the ARID1B NLS mutant using two readouts of canonical Wnt/β-catenin signaling namely a) *AXIN2* expression ^17^ and b) TOP/FOPFlash promoter luciferase assay^10^. As expected, ectopic expression of wild type ARID1B resulted in decreased β-catenin transcriptional activation function (figures 3g-h and S4b). Surprisingly, cytoplasm-restricted ARID1B caused a significant up-regulation of β-catenin transcription activity (figures 3g-h and S4b). β-catenin nuclear entry is inhibited by a destruction complex that triggers proteasomal degradation of cytoplasmic β-catenin which in turn is initiated by phosphorylation at Ser45 position of β-catenin caused by CK1^1^. Indeed, cytoplasm-restricted ARID1B caused an increase in the non-phosphorylated (active) form of β-catenin whereas there was no effect of the wild type ARID1B (figures 3i and S4c).

These observations provided the first probable evidence for a gain of oncogenic function exhibited by cytoplasm-localized ARID1B supporting our initial observations in pancreatic tumor samples.

### Cytoplasm localized ARID1B activates RAF-ERK signaling

We performed a halo tag based pull down-mass spectrometry screen and surprisingly identified components and regulators of the RAF-ERK signaling pathway including a-RAF and c-RAF as potential interactors of cytoplasm-restricted ARID1B (Table S3). RAF-ERK signaling is a major driver of cell viability and growth and is also shown to induce migratory phenotype in epithelial cells ^23^. Of note, previous studies have implicated the RAF-ERK signaling cascade in direct phosphorylation of LRP6 leading to activation of canonical Wnt/β-catenin signaling ^5^ via inhibition of CK1 mediated β-catenin phosphorylation^1^.

We confirmed ARID1B and c-RAF interaction by pull down followed by immunoblotting (figures 4a and S5a) and fluorescence based co-localization studies (figure S5b). More importantly, ectopic expression of cytoplasmic (but not nuclear) version of ARID1B resulted in a significant increase in the levels of active (phosphorylated at Ser338) form of c-RAF and ERK1/2 (figure 4b) though there was no effect on phosphorylated (active) AKT levels (figures 4b and S5c), confirming the specificity of RAF activation by the ARID1B-NLS mutant. Interestingly, wild type ARID1B caused a significant reduction in p-ERK1/2 levels (figure 4b); Wnt/β-catenin signaling is known to activate ERK signaling ^36^ and the observed reduction in active ERK is probably a result of downregulation of β-catenin transcription activation function by wild type ARID1B.

**Figure 4.**
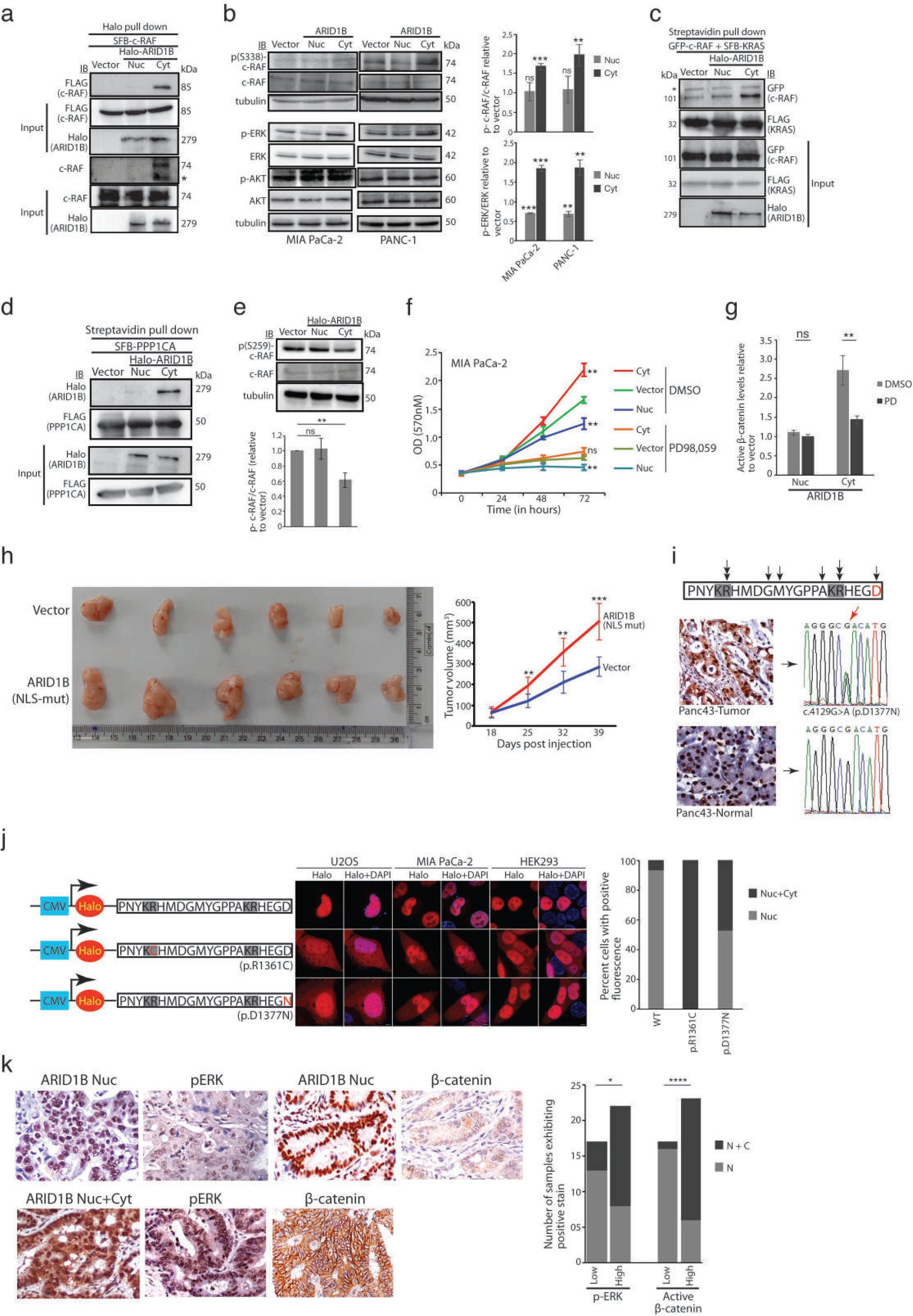
Cytoplasm-localized ARID1B activates RAF-ERK signaling in PaCa cells. **a,** Interaction of ARID1B with both ectopically expressed as well as endogenous c-RAF (to evaluate ARID1B interaction with endogenous c-RAF, transfection with SFB-c-RAF was avoided). **b,** Activation of c-RAF (phosphorylation at Ser 338) and ERK1/2 upon ectopic expression of cytoplasmic form of ARID1B. **c,** Cytoplasm restricted ARID1B increases interaction between constitutively active (mutant) KRAS and c-RAF. The asterisk (*) in panels a and c indicates a non-specific band. **d**, interaction of cytoplasm-localized ARID1B with PPP1CA. **e,** cytoplasm-localized ARID1B decreases levels of phosphorylated (at Ser259) form of c-RAF. **f-g,** Oncogenic function exhibited by cytoplasm-localized ARID1B is dependent on ERK signaling. Results for MTT (panel f; performed as described in figure 3e) and quantitation of active (non-phosphorylated at Ser45) β-catenin levels (panel g; performed as described in figure 3i) upon treatment with the MEK1/2 inhibitor PD98,059 (20μM) are shown. PD, PD98,059. **h,** Oncogenic role of cytoplasm localized ARID1B in mouse xenograft tumor models. A picture of all 12 excised tumors generated by MiaPaCa2 cells expressing cytoplasmic version of ARID1B or vector alone (indicated) is shown in the left. Plot for change in tumor volume (mean for 6 animals) is shown on the right. Error bars represent standard deviation. **i,** Evaluation of ARID1B-NLS mutations. Top panel shows location of eight missense mutations within the ARID1B-NLS identified from the cBioPortal somatic cancer mutation database. Black arrow-heads indicate the amino acids targeted by each mutation; double arrow-heads indicate two distinct mutations affecting the same amino acid. The C-terminal most Aspartic acid residue mutated in pancreatic tumor sample #Panc43 (Table S1) is depicted in red colour. List of mutations is given in Table S4. Bottom panel shows ARID1B immunohistochemistry and DNA sequencing results for tumor and matched normal counterparts of #Panc43 exhibiting cytoplasmic localization of ARID1B. The mutant nucleotide in the electropherogram is indicated by a red arrow-head. **j,** Evaluation of the p.D1377N and p.R1361C ARID1B NLS mutations using the same strategy as in figure 2e. The two basic amino acid clusters are indicted by grey shading and the mutated residue is indicated by red colour. **k**, Oncogenic role of cytoplasm localized ARID1B in human pancreatic tumor samples. ARID1B cytoplasmic localization correlates significantly with active forms of (non-phosphorylated) β-catenin and (phosphorylated) ERK in pancreatic tumor samples. Differences were considered significant at a p value less than 0.05 (*, p<0.05; **, p<0.01; ***, p<0.001; ****, p<0.0001; ns, not significant). Scale bars are 10 µm for all fluorescence images. Nuc, nuclear (wild type) ARID1B; Cyt, cytoplasmic (NLS mutant) ARID1B; IB, immunoblotting. N, nuclear staining; N + C, nuclear plus cytoplasmic staining.

Binding of activated RAS protein to c-RAF is a major inducer of c-RAF activation. We confirmed a significant increase in the interaction between mutant (constitutively activated) KRAS and c-RAF in presence of the cytoplasmic (but not nuclear) form of ARID1B (figure 4c). A critical event facilitating RAS-RAF interaction is the removal of inhibitory phosphorylation at Ser259 residue of c-RAF by protein phosphatases^15^. Interestingly, the list of potential interactors of cytoplasmic ARID1B included PPP1CA (Table S3), the catalytic subunit of a protein phosphatase known to dephosphorylate c-RAF at Ser259^11^. Using the pull down strategy described in figure 4a we confirmed the interaction of cytoplasmic (but not nuclear) ARID1B with PPP1CA (figure 4d). More importantly, the cytoplasmic (but not nuclear) version of ARID1B resulted in decreased levels of phosphorylated (at Ser259) form of c-RAF (figure 4e). Thus, cytoplasmic ARID1B may potentiate removal of inhibitory phosphorylation at Ser259 residue of RAF by promoting the latter’s interaction with PPP1CA providing a possible mechanistic insight into the mode of oncogenic action of cytoplasmic ARID1B.

We repeated the MTT assays (shown in figure 3e) in the presence of PD98,059, a drug that inhibits activity of the MEK1/2 protein kinase responsible for phosphorylation of ERK1/2 downstream of activated RAF. As expected, presence of the blocker reduced overall growth potential of cells but wild type ARID1B was still able to reduce cell viability even in presence of the drug (figure 4f) confirming independence of wild type ARID1B function as well as specificity of PD98,059 action in the assay. More importantly, treatment with the blocker abolished the increased viability of cells caused by the cytoplasmic form of ARID1B (figure 4f) confirming thereby dependency of tumorigenic changes induced by the cytoplasm-localized ARID1B on the activation of ERK signaling pathway. Similarly, the increased levels of *AXIN2* transcript as well as of active (non-phosphorylated) β-catenin observed upon ectopic expression of the cytoplasmic form of ARID1B were significantly reduced in presence of the drug (figures S5d-e). Therefore, the primary effect of cytoplasm localized ARID1B appears to be the activation of RAF-ERK signaling cascade which results in promoting a tumorigenic phenotype besides causing stabilization and activation of β-catenin transcription regulatory function.

### Evidence for oncogenic role of cytoplasmic ARID1B from primary tumor samples and mice tumor xenografts

In order to obtain *in vivo* evidence for possible oncogenic function of cytoplasmic ARID1B, we confirmed significantly increased tumor formation ability of MIA PaCa-2 cells expressing the cytoplasmic form of ARID1B in nude mice xenograft experiments (figures 4h and S6a). We next scrutinized the cBioPortal (cbioportal.org) cancer somatic mutation database and detected several ARID1B mutations including splice site, frame shift and truncations that could potentially inactivate the NLS. More importantly, we identified eight missense mutations located within the NLS (Table S4). Of these, one (p.D1377N) was also detected in a PaCa sample (# Panc43; Table S1) that exhibited significant ARID1B cytoplasmic localization (figure 4i). Two NLS-specific mutations p.R1361C and p.D1377N stimulated cytoplasmic retention of the Halo tag in U2OS, MIA PaCa-2 and HEK293 cells (figures 4j). Finally, IHC based evaluation performed on the PaCa TMA revealed significant correlation between ARID1B cytoplasmic localization and active ERK1/2 and β-catenin levels (figure 4k and Table S5). These results provide support for a possible clinical significance of cytoplasmic localization of ARID1B.

Given that ARID1B is a ubiquitously expressed protein implicated in several cancer types, it’s cytoplasmic localization was unlikely to be restricted to PaCa alone. No cytoplasmic staining was however detected based on IHC performed on TMAs generated for adenocarcinomas of the esophagus (30 tumor and 10 normal samples), colon and rectum (120 tumor and 20 normal samples) as well as squamous carcinomas of the oral tongue (60 tumor and 25 normal samples) and esophagus (80 tumor and 15 normal samples) (data not shown). However, we identified significant ARID1B cytoplasmic localization in breast cancer samples (figures S6b-c and Table S6) suggesting a wider clinical importance of our novel discovery.

## Discussion

In this study, we report a novel oncogenic function for ARID1B emanating from its mis-localization in the cytoplasm. Of the two PaCa cell lines tested, one (PANC-1) harbors the endogenous (nuclear) ARID1B and thus it is possible that the observed effects are due to an interference of the endogenous wild type protein function by the ectopically expressed NLS mutant (similar to a dominant negative effect). However, results obtained with the ARID1B null MIA PaCa-2 cells can only be explained by a gain of function of the cytoplasmic form un-related to its nuclear function. Thus, ARID1B cytoplasmic localization appears to have an additional role to support tumorigenesis not due simply to a loss of its nuclear function.

Our discovery of RAS-RAF-ERK signaling activation through aberrant cytoplasmic localization of ARID1B in pancreatic cancer cells attests to the ‘addiction’ of PaCa cells for the RAS-RAF-ERK signaling cascade as suggested earlier ^32, 35^. Of interest, a synthetic lethality screen in a KRAS mutant cell line revealed ARID1B as a significantly enriched gene ^18^.

Though classified as a tumor suppressor, the SWI/SNF complex ^4, 20, 22^ as well as individual components ^27^ have been shown to exhibit oncogenic functions. Gene fusion events involving the SS18 subunit of the BAF complex exhibiting oncogenic gain of function are reported in synovial sarcomas ^12^. The current study however is probably the first report of mis-localization induced oncogenic gain of function exhibited by a component of the human SWI/SNF complex. We detected ARID1B cytoplasmic localization in a significant proportion of breast cancer samples (classified as ductal adenocarcinoma similar to PaCa) but not in cancers of the colon/rectum, esophagus and the oral tongue.

In conclusion, ARID1B appears to contribute to oncogenesis through non-canonical cytoplasm-based mechanisms in addition to mechanisms involving dysregulation in the nucleus (figure S7), adding to the growing list of cancer genes exhibiting a dual ‘Yin-Yang’ nature^2^. The results presented here have potentially opened an exciting new area of ARID1B and SWI/SNF biology. Is cytoplasmic localization causal to a hitherto undiscovered canonical function of ARID1B? Most tumor types exhibit mono-allelic genetic lesions of SWI/SNF components in contrast to bi-allelic inactivation observed for classical tumor suppressors; the tumor suppression is therefore presumed to be due to a dosage effect and the ‘residual’ SWI/SNF nuclear function has been suggested to be an attractive therapeutic target^19^. One can perhaps envisage a ‘dosage’ effect due to partial ARID1B cytoplasmic localization as well. Given the recent identification of SWI/SNF deficient tumors being susceptible to Bromodomain inhibitors acting through suppression of RAS-ERK signaling^28^ and the recent demonstration of effective pancreatic tumor regression in genetically engineered mice as well as in patient derived mouse xenografts when subjected to combined EGFR/c-RAF ablation^3^, the novel RAF-ERK activating function of cytoplasm-localized ARID1B can be explored for therapeutic options.

## Materials and methods

### Construction of tissue microarray (TMA), immunohistochemistry (IHC), microdissection and DNA sequencing

The PaCa TMA described earlier ^14^ was expanded by the addition of fifteen samples to contain a total of 67 tumor and matched normal sample pairs. A Breast cancer TMA was constructed in a similar way and included 65 tumor and 30 normal samples. Details of sample collection and TMA construction are provided in supplementary methods. ARID1B ^14^, β-catenin ^24^ and pERK IHC were performed using standard protocol on 4 µM sections; details are given in supplementary methods section. For ARID1B, cores exhibiting ≥20% (nuclear or cytoplasmic) epithelial staining were classified as positive. For β-catenin, scores for stain intensity (negative (0), weak (1), moderate (2) and strong (3)) were added with those for percentage staining and summated scores of 1-3 and 4-7 were considered low and high expression, respectively. For pERK, the percent epithelial staining was converted to a numerical score (up to 25 (1), 50 (2), 75 (3) and 100% (4)) and classified as negative/low (1-2) or high (3-4) expression. Images were taken using Nikon eclipse 80i (Nikon corporations) at 20X magnification. All antibodies are listed in supplementary methods.

DNA was extracted from tumor epithelium micro-dissected from Formalin Fixed Paraffin Embedded (FFPE) sections of PaCa samples by using the standard Phenol-Chloroform / ethanol method ^24^. The NLS encoding region was amplified using specific primers at an annealing temperature of 55^0^C using Amplitaq Gold^TM^ (ABI Inc.). The PCR product was sequenced using the 3100 genetic analyzer (ABI Inc.).

### Cell line maintenance and their manipulations

MIA PaCa-2 and PANC-1 cells were procured from ATCC while U2OS, and HEK293 cells were kind gifts from Dr. Rashna Bhandari and Dr. Sangita Mukhopadhyay (CDFD, Hyderabad, India), respectively. Details of maintenance of cell lines, transfection and various assays are provided in the supplementary methods section.

### Pull down, immunoblotting and Mass spectrometry

Halo tag (Halo pulldown and labeling kit, Promega Corporations) and Streptavidin bead (GE Health care Bio-Science) based pull down were performed as per standard protocol and the elutes were resolved on SDS-PAGE, transferred onto PVDF membrane (Merck Millipore) and probed with primary antibody at 4^0^C overnight. Immuno-reactive bands were detected following treatment with secondary antibodies using ECL™ Prime Western Blotting detection reagent. Mass spectrometry analysis was outsourced to the Taplin Biological Mass Spectrometry Facility (Harvard Medical School). All antibodies are listed in supplemental methods.

### Nude mice xenograft experiments

All nude mice experiments were performed in accordance with the institute guidelines and were approved by the Institutional Animal Ethics Committee. 8-9 weeks old female FOXN1^-/-^ nude mice (n=6), were subcutaneously injected with stable MIA PaCa2 cells (5.0 × 10^6^) expressing either cytoplasmic form of ARID1B or empty vector. Tumor diameters were serially measured with a digital calliper every 7 days and tumour volumes were calculated using the formula: Volume=(W^2^*L)/2, where W and L represent width and length respectively. Mice were sacrificed 6 weeks post injection by carbon dioxide euthanasia, tumors were dissected and weight and volume were measured.

### Statistical analysis

All data obtained from three independent experiments (biological replicates) were represented as mean +/-standard deviation. Student’s t test was used to determine the statistical significance for all experiments while the Fisher exact test (two-tailed) was used to calculate statistical significance of IHC data.

## Acknowledgements

The ARID1B cDNA construct was a kind gift from Dr Reiko Watanabe, Tohoku University, Sendai City, Japan. Antibodies against c-RAF and (Ser 338)p-c-RAF were generous gifts from Dr Atin Mandal, Bose Institute, Kolkatta, India. pDEST-SFB-PPP1CA, pDEST-N-SFB, pDEST-N-GFP and TOP/FOPFlash vectors were kind gifts from Dr MS Reddy, CDFD, Hyderabad, India. The mutant (G12V) KRAS cDNA construct was a kind gift from Dr. S K Manna, CDFD, Hyderabad, India. U20S and HEK293 cell lines were kind gifts from Dr. Rashna Bhandari, CDFD, Hyderabad, India and Dr. Sangita Mukhopadhyay, CDFD, Hyderabad, India, respectively. The work was supported by a grant (BT/PR13948/BRB/10/1406/2015) from the Department of Biotechnology, Government of India to MDB. SA, a registered PhD student of Manipal Academy of Higher Education, is grateful to the Department of Science and Technology, Government of India for junior and senior research fellowships. We acknowledge CDFD’s Sophisticated Equipment Facility for fluorescence microscopy and Sanger sequencing and the Experimental Animal Facility, for nude mice experiments. We thank the patients for kindly agreeing to be a part of the study. We acknowledge Dr C Sundaram and Dr Shantveer Uppin, Nizam’s Institute of Medical Sciences, Hyderabad for providing Formalin Fixed Paraffin Embedded (FFPE) blocks of PaCa samples. We are grateful to Dr. J Gowrishankar, CDFD, Hyderabad, India and Dr Geeta Narlikar, UCSF, San Francisco, CA, USA, for critical reading and important suggestions on the manuscript.

## Author contributions

MDB supervised the research, arranged funding and wrote the manuscript. MDB and SA conceived the project, designed the experiments and analyzed the results. SA, performed experiments involving fluorescence, functional assays and immunoblotting and contributed to manuscript writing. SA and PK performed cloning and Q-PCR experiments. PK performed luciferase assays. Experiments related to construction of the TMA, IHC and mutation screening from tumor samples were performed by VK. All authors contributed to manuscript correction.

## Supplementary Figure Legends

**Figure S1. IHC-based comparative assessment of ARID1B expression in pancreatic tumor and normal samples** (total fifty five tumor/normal pairs). Differences were considered significant at a p value less than 0.05 (***, p<0.001).

**Figure S2. Clustal Omega based analysis of the ARID1B-NLS. a,** Homology between ARID1B protein sequence of 42 metazoan species. The alignment is shown only for the region comprising the putative human ARID1B NLS. Symbols used to indicate amino acid conservation are ‘*’, identity; ‘:’, conserved substitution; ‘.’, semi-conserved substitution. The two highly conserved basic amino acid clusters (Lysine-Arginine) are indicated by a thick red bar on the top. Identities of the forty two species used for Clustal Omega alignment are *Homo sapiens*, *Mus musculus, Rattus norvegicus*, *Danio rerio*, *Bos taurus*, *Myotis lucifugus*, *Ornithorhynchus anatinus*, *Ictidomys tridecemlineatus*, *Ailuropoda melanoleuca*, *Ovis aries*, *Macaca mulatta*, *Sus scrofa*, *Canis lupus familiaris*, *Pan troglodytes*, *Pongo abelii*, *Sarcophilus harrisii*, *Taeniopygia guttata*, *Callithrix jacchus*, *Mustela putorius furo*, *Felis catus*, *Ficedula albicollis*, *Gallus gallus*, *Anas platyrhynchos*, *Anolis carolinensis*, *Equus caballus*, *Monodelphis domestica*, *Loxodonta africana*, *Oreochromis niloticus*, *Meleagris gallopavo*, *Gorilla gorilla gorilla*, *Nomascus leucogenys*, *Oryctolagus cuniculus*, *Papio anubis*, *Xenopus tropicalis*, *Ophiophagus hannah*, *Xiphophorus maculates*, *Chlorocebus sabaeus*, *Astyanax mexicanus*, *Poecilia formosa*, *Tetraodon nigroviridis*, *Takifugu rubripes* and *Alligator mississippiensis*. **b,** Alignment of ARID1B and ARID1A protein sequence reveals significant conservation of their respective NLS sequences. Figure shows alignment of only the NLS region; the N- and C-terminal basic amino acid clusters are highlighted by grey shading. Symbols used to indicate amino acid conservation are ‘*’, identity and ‘:’, conserved substitution.

**Figure S3. Characterization of the ARID1B-NLS. a,** Importazole treatment blocks ARID1B nuclear localization in HEK293 cells. Scale bars are 10µm. **b,** The ARID1B NLS peptide can translocate a fused GFP tag into the nucleus. Basic amino acid clusters are indicated by grey shading while the mutant residues are shown in red color. Representative images obtained separately from cells transfected with each construct are shown. Scale bars are 10µm for U2OS and HEK293 and 5µm for MIA PaCa-2.

**Figure S4. Cytoplasmic localization due to mutant NLS sequence compromises ARID1B canonical functions. a,** Cytoplasm-restricted ARID1B is compromised in its ability to activate *CDKN1A/B* in MIA PaCa-2 cells as evaluated through *CDKN1A/B* promoter luciferase assays. **b-c,** Cytoplasm-restricted ARID1B enhances β-catenin transcription activity in PANC-1 cells. **b,** Quantitation of *AXIN2* relative mRNA levels. **c,** Quantitation of active (non-phosphorylated at Ser45) β-catenin levels. Nuc, nuclear (wild type) ARID1B; Cyt, cytoplasmic (NLS mutant) ARID1B. Differences were considered significant at a p value less than 0.05 (*, p<0.05; **, p<0.01; ***, p<0.001; ns, not significant).

**Figure S5. Characterization of oncogenic role of cytoplasm-localized ARID1B. a,** ARID1B and c-RAF interaction was analyzed using pull down followed by immunoblotting. **b,** ARID1B-c-RAF interaction is shown using fluorescence based co-localization. White arrowheads indicate sites of co-localization of cytoplasm localized ARID1B with c-RAF. Scale bars are 10µm. **c,** Cytoplasm-localized ARID1B causes no significant change in levels of active AKT (p-AKT). Figure shows results of densitometric quantitation of active/total ratios of AKT (from figure 4b) represented as fold induction over the levels obtained with the pDEST-N-EGFP vector. **d-e,** Oncogenic function exhibited by cytoplasm-localized ARID1B is dependent on ERK signaling. Results for *AXIN2* mRNA induction (panel d; performed as described in figure 3g) and quantitation of active (non-phosphorylated at Ser45) β-catenin levels (panel e; performed as described in figure 3i) upon treatment with the MEK1/2 inhibitor PD98,059 (20μM) are shown. IB, immunoblotting. Nuc, nuclear (wild type) ARID1B; Cyt, cytoplasmic (NLS mutant) ARID1B. Differences were considered significant at a p value less than 0.05 (*, p<0.05; **, p<0.01; ns, not significant).

**Figure S6. Assessment of oncogenic potential of cytoplasmic ARID1B in mouse tumor models and human primary breast tumors. a,** Left panel shows picture of all six mice used for xenograft assays following euthanization. Plots for final tumor volume and weight (after excision) are shown on the right. **b,** representative IHC images depicting ARID1B localization in primary breast tumors. **b,** comparative analysis of number of samples exhibiting ARID1B nuclear vs nuclear + cytoplasmic stain in tumor and normal samples (fifty eight tumor and thirty normal samples). Differences were considered significant at a p value less than 0.05 (**, p<0.01; ***, p<0.001). N, nuclear staining; N + C, nuclear plus cytoplasmic staining.

**Figure S7. A model depicting contrasting roles for nuclear and cytoplasmic localized ARID1B based on results obtained from this study.**

